# Modeling disease-correlated TUBA1A mutation in budding yeast reveals a molecular basis for tubulin dysfunction

**DOI:** 10.1101/2020.04.13.039982

**Authors:** E. Denarier, K.H. Ecklund, G. Berthier, A. Favier, S. Gory, L. De Macedo, C. Delphin, A. Andrieux, S.M. Markus, C. Boscheron

## Abstract

Malformations of cortical development (MCD) of the human brain are a likely consequence of defective neuronal migration, and/or proliferation of neuronal progenitor cells, both of which are dictated in part by microtubule-dependent transport of various cargoes, including the mitotic spindle. Throughout the evolutionary spectrum, proper spindle positioning depends on cortically anchored dynein motors that exert forces on astral microtubules emanating from spindle poles. A single heterozygous amino acid change, G436R, in the conserved TUBA1A α-tubulin gene was reported to account for MCD in patients. The mechanism by which this mutation disrupts microtubule function in the developing cerebral cortex is not understood. Studying the consequence of tubulin mutations in mammalian cells is challenging partly because of the large number of α-tubulin isotypes expressed. To overcome this challenge, we have generated a budding yeast strain expressing the mutated tubulin (Tub1^G437R^ in yeast) as one of the main sources of α-tubulin (in addition to Tub3, another α-tubulin isotype in this organism). Although viability of the yeast was unimpaired by this mutation, they became reliant on Tub3, as was apparent by the synthetic lethality of this mutant in combination with *tub3*Δ. We find that Tub1^G437R^ assembles into microtubules that support normal G1 activity, but lead to enhanced dynein-dependent nuclear migration phenotypes during G2/M, and a consequential disruption of spindle positioning. We find that this mutation impairs the interaction between She1 – a negative regulator of dynein – and microtubules, as was apparent from a yeast two-hybrid assay, a co-sedimentation assay, and from live cell imaging. We conclude that a weaker interaction between She1 and Tub1^G437R^-containing microtubules results in enhanced dynein activity, ultimately leading to the spindle positioning defect. Our results provide the first evidence of an impaired interaction between microtubules and a dynein regulator as a consequence of a tubulin mutation, and sheds light on a mechanism that may be causative of neurodevelopmental diseases.

## Introduction

Malformations of cortical development (MCD) are severe brain malformations associated with intellectual disability and infantile refractory epilepsy. MCD include lissencephaly, pachygyri, and polymicrogyria, brain malformations characterized by alterations in cortical gyration and sulcation. These diseases have a strong genetic basis involving the α- and β-tubulin subunits of microtubules, as well as various microtubule associated proteins (MAPs) and effectors. For example, mutations in LIS1 – an effector of microtubule based transport – account for ∼65% of classic lissencephaly (Kumar et al., 2010). LIS1 affects microtubule-based transport by activating the motility of the molecular motor cytoplasmic dynein-1 (Marzo et al., 2020; Mohamed M. Elshenawy, 2020; Zaw Min Htet, 2020), which is also found mutated in MCD patients (Laquerriere et al., 2017; Poirier et al., 2013; Vissers et al., 2010; Willemsen et al., 2012). Recently, a patient with pachygyria and severe microcephaly associated with postural delay and poor communication abilities was shown to possess a single *de novo* heterozygous mutation in the α-tubulin gene, TUBA1A, that results in a glycine to arginine substitution at position 437 (Bahi-Buisson et al., 2008). Although currently unclear, the cellular basis for disease in this patient may be defects in mitotic spindle orientation, which is a critical process during asymmetric cell division that is required for cellular differentiation and determination of daughter cell fate (Cabernard and Doe, 2009; Das and Storey, 2012; Williams et al., 2011; Wu et al., 2010). Defects in this process can limit the number of progenitor cells, and ultimately the neuronal mass of the developed brain (Bershteyn et al., 2017). This is apparent in mouse neuronal progenitor cells in which dynein dysregulation by Lis1 inhibition impairs microtubule dynamics, results in a mitotic spindle mispositioning phenotype, and leads to a consequential decrease in the number of progenitors during early development (Tsai et al., 2005; Yingling et al., 2008).

We hypothesized that the TUBA1A G436R mutation might cause disease by leading to defective spindle positioning. This may be due to: (1) alteration of microtubule dynamics and/or structure; or, (2) impairment of motor or MAP function due to disrupted microtubule binding. Distinguishing between these possibilities and deciphering the precise molecular defects arising from tubulin mutations is not a trivial task. Studying tubulin mutations in mammalian cells is complicated by the fact that numerous isoforms of α and β-tubulin are present in the human genome (9 α-tubulin, and 10 β-tubulin isoforms) (Findeisen et al., 2014; Khodiyar et al., 2007). Each of these has a distinct expression pattern, and thus every cell’s tubulin content is a composite mixture of these many variants. In contrast to higher eukaryotes, things are much simpler in the budding yeast *Saccharomyces cerevisiae* in which ∼70-90% of α-tubulin is expressed from the essential *TUB1* gene, with the remaining ∼10-30% arising from the *TUB3* gene (Bode et al., 2003; Gartz Hanson et al., 2016; Schatz et al., 1986b). In addition to its simplicity, the mechanisms and effectors of spindle orientation (e.g., dynein, Lis1) and microtubule dynamics and function (*e.g.*, tubulin, EB1, CLIP-170, ChTOG) are highly conserved between humans and budding yeast. In this organism, it is mandatory that the mitotic spindle is correctly positioned along the mother-daughter cell axis and in close proximity to the bud neck prior to mitotic exit, otherwise cell viability is compromised. The reliance on this process for viability permitted the use of genetic screens that revealed the presence of two distinct pathways that can effect this process: namely, the Kar9/actomyosin and dynein pathways (Miller and Rose, 1998). The Kar9/actomyosin pathway relies on a microtubule guidance mechanism, whereby a microtubule plus end-associated myosin (Myo2) orients the mitotic spindle along the mother-daughter axis (Hwang et al., 2003; Lee et al., 2000; Yin et al., 2000). Myo2 is recruited to microtubule plus ends by the concerted effort of Kar9 (homolog of human adenomatous polyposis coli tumor suppressor, APC) and the autonomous microtubule plus end-tracking protein, Bim1 (homolog of human EB1). Recently, we have modeled the disease-correlated TUBB2B F265L β-tubulin mutation in budding yeast, and found that this mutation specifically compromised the Kar9/actomyosin pathway by disrupting the plus end localization of Bim1 (Denarier et al., 2019).

Cytoplasmic dynein, on the other hand, functions from the cell cortex, from where Num1-anchored motors walk along microtubules emanating from spindle pole bodies (the equivalent of centrosomes), which results in the positioning of the spindle at the mother-bud neck (Carminati and Stearns, 1997; Heil-Chapdelaine et al., 2000; Li et al., 1993). Dynein is delivered to Num1 receptor sites at the bud cortex by a two-step “offloading” mechanism: (1) microtubule plus end-associated Bik1 (homolog of human CLIP170) recruits dynein-Pac1 (homolog of human Lis1) complexes to dynamic plus ends (Badin-Larcon et al., 2004; Caudron et al., 2008; Lee et al., 2003; Sheeman et al., 2003); (2) plus end-associated dynein, which appears to be inactive (Lammers and Markus, 2015) is delivered, or “offloaded” to cortical Num1 receptor sites along with its effector complex, dynactin (Markus and Lee, 2011). The extent of dynein activity is largely governed by its localization to these sites; however, as in higher eukaryotes (Tan et al., 2019), at least one known MAP can also regulate dynein activity in cells: She1. The precise mechanism by which it does so in cells is currently unclear; however, *in vitro* studies show that She1 can reduce dynein velocity through simultaneous interactions with both microtubules and dynein (Ecklund et al., 2017), whereas live cell studies have shown that She1 plays a role in polarizing dynein-mediated spindle movements toward the daughter cell (Markus et al., 2012), perhaps in part by tuning dynactin recruitment to plus end-associated dynein (Markus et al., 2011; Woodruff et al., 2009).

To gain insight into the role and importance of α-tubulin G436 (hereafter referred to as G437, due to its position in yeast α-tubulin), and how it might affect the above processes, we produced *S. cerevisiae* yeast strains in which the native *TUB1* locus was replaced with the G437R mutant allele (*tub1*^*G437R*^). Our results show that this mutation leads to alterations in microtubule dynamics, and a spindle positioning defect that is likely due to dysregulated dynein function. The dynein dysfunction phenotype is not a consequence of its mislocalization, but is more likely due to a reduced association of She1 with the mutant microtubules. Although there is no clear She1 homolog in human cells, we propose that this mutation might similarly interfere with dynein function by disrupting the microtubule binding behavior of a regulatory MAP, thus leading to neuronal physiological deficits, and a consequent disruption of cerebral cortex development.

## Results

### *TUB1 G437R* mutagenesis leads to an enhancement in microtubule dynamics during G2/M phase

Glycine 436 of TUBA1A α-tubulin, mutation of which is highly correlated with a developmental human brain disease, is conserved among α-tubulins from numerous organisms, including budding yeast (*TUB1*; overall 74.1% identity between human TUBA1A and yeast Tub1). Glycine 436 (Fig. 1A, red sphere) is one of three highly conserved small, hydrophobic residues in α-tubulin (Fig. 1B, red box) that immediately precede the disordered carboxy-terminal tail of α-tubulin. This region of α-tubulin partly constitutes the external surface of the microtubule to which MAPs and motor proteins bind (Fig. S1). Although some MAPs (*i.e.*, Tau and Tpx2) bind proximal to G437, this residue does not appear to encompass the kinesin and dynein binding interfaces (see Fig. S1) (Alushin et al., 2014; Lowe et al., 2001; Nogales et al., 1998). Modeling an arginine into position 436 of porcine α-tubulin (pdb 3J6G; using UCSF Chimera (Pettersen et al., 2004)) revealed that such a mutation may potentially disrupt the terminal helix of α-tubulin (note the steric clash between the modeled arginine and helix 11 in rotational isomers 1 and 4; Fig. S2). To assess the phenotypic consequences of this mutation, we engineered yeast strains to express Tub1^G437R^, in which the mutation was introduced at the native *TUB1* locus.

**Figure 1.**
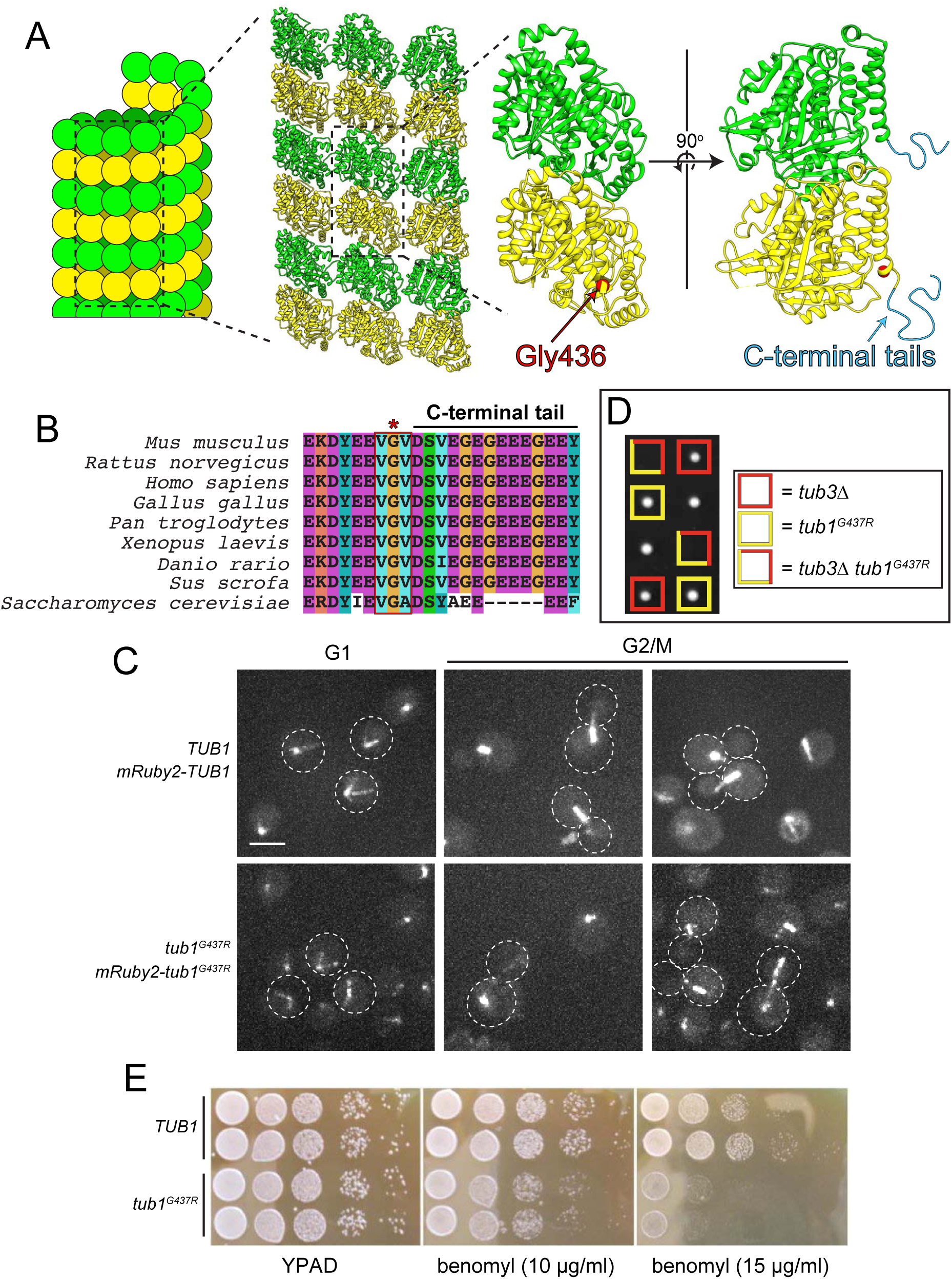
The α-tubulin G437R mutant is a polymerization competent tubulin in yeast. (A) Structural model of the microtubule lattice with Gly436 indicated (red circle). A cartoon of the unstructured C-terminal tail (in cyan) is included on the right (the model is derived from PDB 3J6G (Alushin et al., 2014)). (B) Alignment of the C-terminal region of α-tubulin from various species. The red box delineates a conserved region of small, hydrophobic residues that immediately precedes the C-terminal tail, and the red asterisk indicates Gly436. Amino acids are colored according to the Clustal color scheme. (C) Representative fluorescence images of yeast cells with indicated genotype at different stages of their cell cycle (as determined from cell and spindle morphology). Scale bar, 5 µm. (D) Representative image of colonies (grown on YPAD) from a tetrad dissection depicting synthetic lethal interaction between *tub1*^*G437R*^ and *tub3*Δ. Two representative tetrads are shown. (E) Growth assay with haploid cells of indicated genotype incubated on rich media (YPAD) with or without the indicated concentration of benomyl, a microtubule destabilizing drug.

Heterozygous diploid cells (*TUB1*/*tub1*^*G437R*^) were sporulated, and the resulting haploid *tub1*^*G437R*^ mutant cells were recovered at the expected frequency. The mutants exhibited no growth defects when cultured in nutrient-rich media (YPAD; see Fig. 1E) indicating that the mutation does not compromise yeast cell viability. Using a chromosomally-integrated *RFP-tub1*^*G437R*^ (expressed in the presence of the untagged *tub1*^*G437R*^ allele), we found that the mutant tubulin incorporates into spindle and cytoplasmic microtubules during all phases of the yeast cell cycle (G1, G2/M; Fig 1C).

In addition to *TUB1*, budding yeast possess a second α-tubulin gene encoded by *TUB3*. Whereas *TUB1* is the major α-tubulin isotype and is required for cell viability, cells tolerate deletion of *TUB3*, which possesses 91% identity and 95% similarity with *TUB1* (Schatz et al., 1986a). To determine if cells could tolerate expressing only Tub1^G437R^, we generated heterozygous *TUB1*/*tub1*^*G437R*^ *TUB3*/*tub3Δ* diploid cells, sporulated them, and assessed viability of the recovered single and double mutant progeny. Although the single mutants exhibited relatively normal colony morphology, none of the double mutants were viable (6 out of 6 expected double mutants were inviable), revealing that expression of only the mutant G437R α-tubulin leads to cell death (Fig. 1D). Although the reason for the inviability of the double mutants is unclear, it suggests that either microtubules may not be assembled from only the mutant α-tubulin protein, or microtubules assembled from only the mutant α-tubulin protein do not support proper function. As a consequence of the inviability of the double mutants, we focused the remainder of our study on the *tub1*^*G437R*^ single mutants (*i.e.*, in the presence of wild-type *TUB3*).

Increased sensitivity of cells to the microtubule depolymerizing drug benomyl is a common phenotype of strains with α-tubulin and β-tubulin mutations (Richards et al., 2000). We assessed benomyl sensitivity of wild-type (*TUB1*) and *tub1*^*G437R*^ cells by spotting a dilution series of each on solid media containing different concentrations of the drug (10 and 15 μg/ml; Fig. 1E) and examining cell growth. In the presence of benomyl, cell growth was markedly impaired for *tub1*^*G437R*^ cells compared to wild-type cells, indicating an enhanced sensitivity to the drug as a consequence of the mutant tubulin. This suggests that the G437R mutant could be altering microtubule stability or dynamics.

To determine if this was the case, we measured microtubule dynamics parameters by tracking the movement of microtubule plus ends with Bik1-GFP, the homolog of human CLIP-170 (see Fig. 3A). While we did not observe any notable differences in microtubules dynamics parameters between wild-type and *tub1*^*G437R*^ strains in G1 cells, we did note several differences during the G2/M phase of the cell cycle (Fig. 2). In particular, we noted an increase in the rates of polymerization (1.4 µm/min in wild-type, versus 1.7 µm/min in mutant cells) and depolymerization for microtubules in *tub1*^*G437R*^ cells (1.6 µm/min versus 2.4 µm/min). We also observed a significant increase in the fraction of time the microtubules spent in their growth phase, and a concomitant reduction in the relative fraction of time spent in pause in the mutant cells. This resulted in an overall increase in microtubule dynamicity (Toso et al., 1993) (Fig. 2), which may account for the enhanced sensitivity of the mutant cells to the depolymerizing agent, benomyl. Finally, although the mean microtubule length did not significantly differ between the two strains (Fig. S3A), we noted that the mutant cells exhibited a larger fraction of long microtubules (38% of microtubules were ≥ 7µm in *tub1*^*G437R*^ cells, versus 10% in wild-type cells; Fig. S3B and C), many of which extended from one cell compartment to the other (also see below).

**Figure 2.**
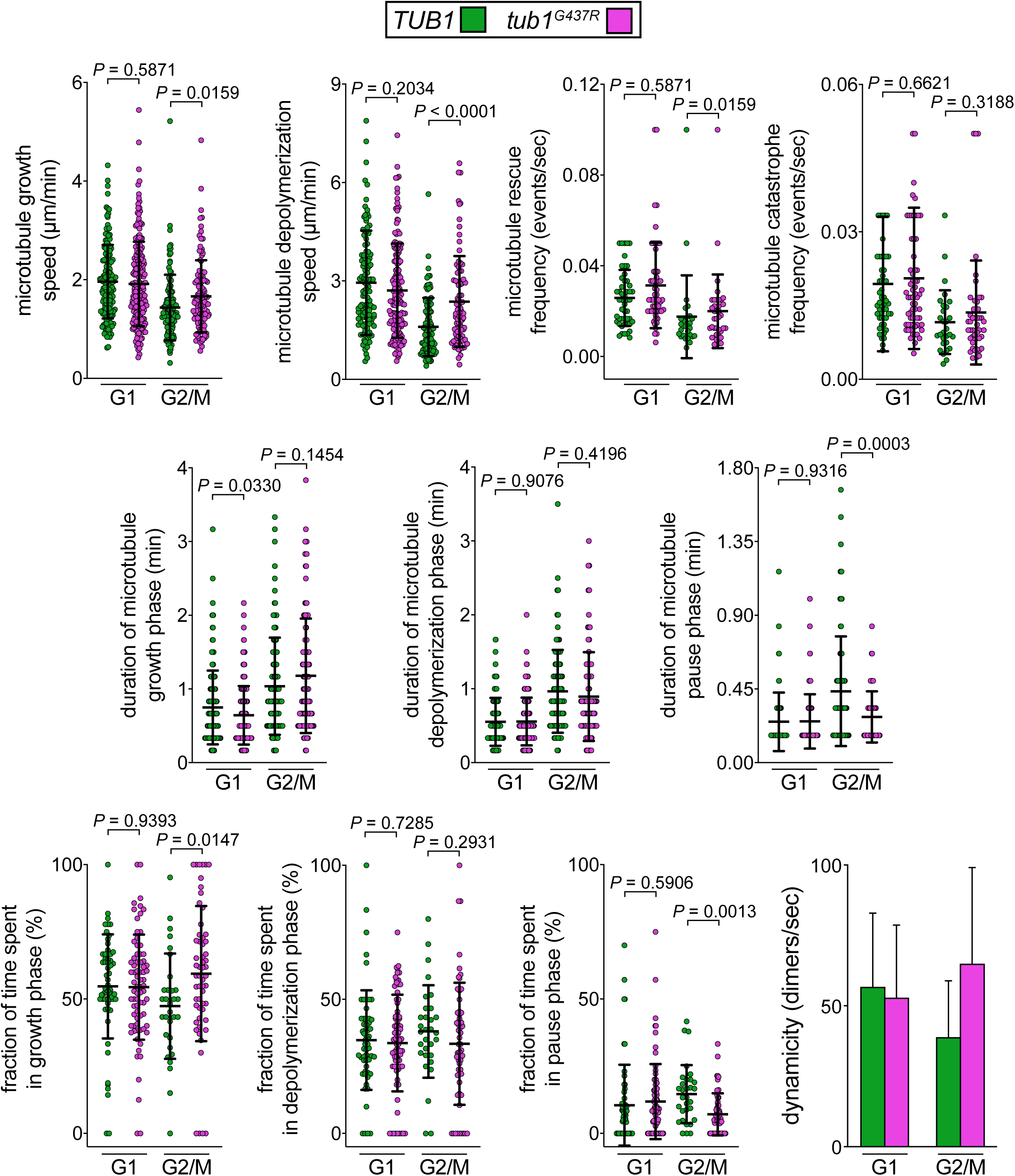
*tub1*^*G437R*^ cells exhibit altered microtubule dynamics during G2/M phase. Plots depicting the indicated microtubule dynamics parameters in the indicated phase of the cell cycle (as determined by cell and spindle morphology). Microtubule behavior was tracked over time using Bik1-GFP as a reporter (localizes prominently to the spindle, to varying degrees along microtubules, and at microtubule plus ends). With the exception of the dynamicity plot (showing mean values with standard deviations), all plots depict all data points (scatter plots) along with mean and standard deviations (bars). *P* values were calculated using an unpaired Welch’s t test.

**Figure 3.**
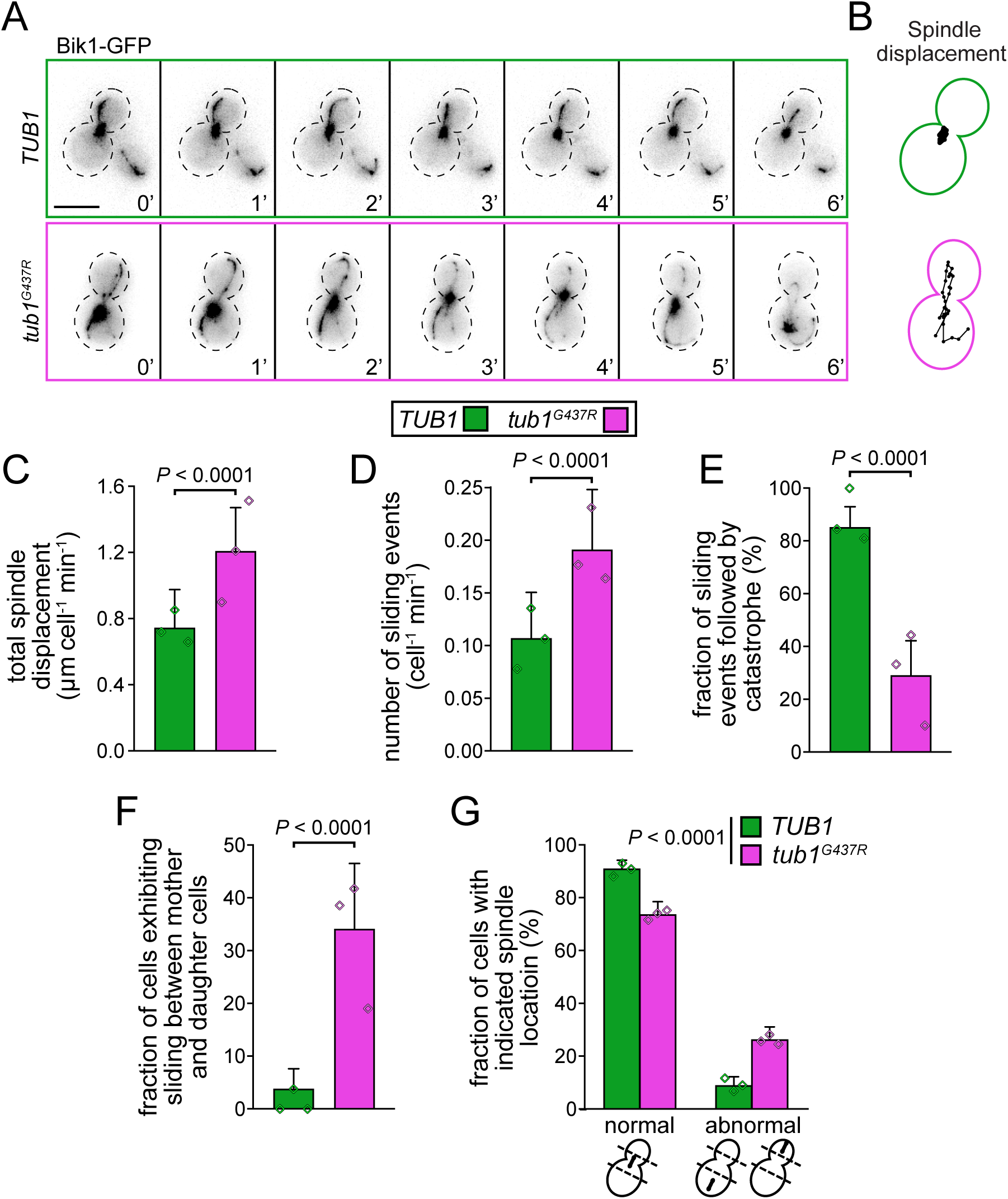
Tub1^G437R^ leads to enhanced spindle dynamics and a spindle misorientation phenotype. (A) Representative inverse fluorescence time-lapse images of wild-type (*TUB1*) and *tub1*^*G437R*^ mutant cells expressing Bik1-GFP. Note the increased spindle movements that are coincident with microtubule sliding events in the mutant cell (bottom; see 3’ – 6’ in the *tub1*^*G437R*^ mutant example for a typical sliding event). Scale bar, 5 µm. (B) Displacement traces of the spindle noted over the 10 min acquisition period (each spot represents the position of the spindle centroid over time). (C and D) Plots depicting the average total displacement of the mitotic spindles in wild-type or mutant cells per minute of the acquisition period (C; n = 74, and 52 cells from three independent clones for *TUB1* and *tub1*^*G437R*^ strains, respectively), or the number of microtubule sliding events (in which a microtubule appears to slide along the cell cortex) per cell per minutes (D; n = 57, and 41 cells from three independent clones, for *TUB1* and *tub1*^*G437R*^ strains, respectively). Bars represent mean values with standard error. (E) Fraction of microtubule sliding events (described above) that are followed by a microtubule catastrophe (n = 40, and 31 cells from three independent clones, for *TUB1* and *tub1*^*G437R*^ strains, respectively). Bars represent weighted means with weighted standard error of proportion. (F) Fraction of cells exhibiting microtubule sliding events in which a very long microtubule (extending from one cell compartment to the other) slides over the bud and mother cell prior to terminating (n = 57, and 46 cells from three independent clones, for *TUB1* and *tub1*^*G437R*^ strains, respectively). Bars represent weighted means with weighted standard error of proportion. (G) Plot depicting the fraction of cells with the indicated spindle location in control and mutant cells (n = 233, and 255 cells from three independent clones, for *TUB1* and *tub1*^*G437R*^ strains, respectively. Bars represent weighted means with weighted standard error of proportion. For panels C and, D, *P* values were calculated using an unpaired Welch’s t test; for panels E – G, statistical significance was determined by calculating Z scores as previously described (Marzo et al., 2019). For all plots, diamonds represent mean values obtained from each independent replicate experiment.

### The G437R mutation leads to increased spindle dynamics and impaired spindle positioning

Since the G437R mutation specifically affects microtubule dynamics during G2/M phase, during which astral microtubules effect mitotic spindle movements, we sought to assess the consequence of mutagenesis on mitotic spindle dynamics during this phase using Bik1-GFP as a fluorescent reporter (Fig. 3A). In wild-type cells, spindles sampled a relatively small area near the bud neck in the mother cell, and the majority of them were oriented along the mother-bud axis (Fig. S4, and Fig. 3A and B, top). Although the majority of spindles in *tub1*^*G437R*^ cells were also oriented along the mother-bud axis (Fig. S4), they exhibited highly dynamic behavior; specifically, we observed numerous instances of the spindle oscillating back and forth between the mother and daughter cell compartments (Fig. 3A and B, bottom). To quantitate this phenomenon, we measured the total distance over which the mitotic spindle moved per minute within each cell. Compared with wild-type cells (*TUB1*), the spindles in *tub1*^*G437R*^ cells moved a significantly longer distance (0.7 µm versus 1.2 µm; Fig. 3C). These observations indicate that excessive forces emanating from the mother and bud cortex are exerted upon the mitotic spindle in *tub1*^*G437R*^ cells. We also noted that the spindle movements in *tub1*^*G437R*^ cells occurred coincidently with microtubule “sliding” events, during which the plus end of the astral microtubule contacts the cell cortex and then curls along it, all the while maintaining lateral contact (see Fig. 3A, bottom, 3’ thru 6’; also see Video S1). Quantitation of these movements – which are characteristic of dynein-mediated spindle movement events (Adames and Cooper, 2000) – revealed an approximate two-fold increase in their frequency in *tub1*^*G437R*^ cells with respect to wild-type cells (Fig. 3D).

A recent study found that dynein-mediated microtubule sliding events are often followed by a microtubule catastrophe (also mediated by dynein), which plays a role in attenuating the spindle movement events (Estrem et al., 2017). We noted that these particular microtubule catastrophe events (*i.e.*, those following sliding events) are greatly reduced in *tub1*^*G437R*^ cells (29% of sliding events are followed by catastrophe in *tub1*^*G437R*^ cells, versus 85% in wild-type cells; Fig. 3E), suggesting that the G437R mutation reduces dynein’s ability to induce a catastrophe. Also consistent with these data, a larger fraction of mutant cells appeared to exhibit events in which very long microtubules that extended from one compartment to the other (*i.e.*, mother to daughter, or vice versa) underwent characteristic dynein-mediated sliding between the two cellular compartments (Fig. 3F; also see Video S1).

The main function of cytoplasmic microtubules in vegetative yeast cells is to orient the mitotic spindle along the mother-daughter axis (by the Kar9/actomyosin pathway), and localize it proximally to the bud neck (by the dynein pathway) such that at the time of anaphase onset, the chromosomes are divided equally between the mother and daughter cells (Carminati and Stearns, 1997; Hwang et al., 2003; Li et al., 1993; Liakopoulos et al., 2003; Yin et al., 2000). Although we found that orientation of the spindle along the mother-daughter axis was not compromised in *tub1*^*G437R*^ cells (Fig. S4) – suggesting the Kar9/actomyosin pathway is not compromised – we did note that the spindle was more frequently localized to the apical regions of the mother or daughter cells in the *tub1*^*G437R*^ strain (26% of *tub1*^*G437R*^ cells, versus 9% of wild-type cells; Fig. 3G). Taken together, these observations suggest that the G437R mutation leads to increased dynein-mediated spindle movements, and yet reduced dynein-mediated microtubule catastrophe events (following sliding events), which ultimately leads to a spindle mislocalization phenotype.

### Enhanced spindle dynamics in *tub1*^*G437R*^ mutant cells are dynein-dependent

As noted above, the dynein pathway effects microtubule “sliding” events that result in translocation of the mitotic spindle throughout the cell. Given our observations noted above, we sought to determine whether the *tub1*^*G437R*^ mutant phenotypes are a consequence of hyperactive dynein. Consistent with the notion that dynein is responsible for the observed spindle behavior in *tub1*^*G437R*^ cells, the increased spindle displacement phenotype in *tub1*^*G437R*^ cells was eliminated by deletion of *DYN1* (which encodes for the dynein heavy chain), but not by deletion of *KIP3*, a kinesin that has been implicated in regulating microtubule length and spindle movements (Fukuda et al., 2014; Gupta et al., 2006).

We next asked whether the increased dynein activity is due to enhanced targeting of dynein to microtubule plus ends or the cell cortex, which can be causative of increased cellular dynein activity (Markus et al., 2011). To this end, we imaged Dyn1-3GFP in cells also expressing RFP-Tub1 (or RFP-Tub1^G437R^) in wild-type or *tub1*^*G437R*^ cells (Fig. 4C - E). We found no significant difference in either the frequency of Dyn1-3GFP targeting, or the fluorescence intensity values for Dyn1-3GFP at microtubule plus ends, spindle pole bodies (SPBs; equivalent of centrosomes), or the cell cortex (Fig. 4D and E). Thus, the increased dynein activity in *tub1*^*G437R*^ cells is likely not a consequence of increased localization to any of these sites.

**Figure 4.**
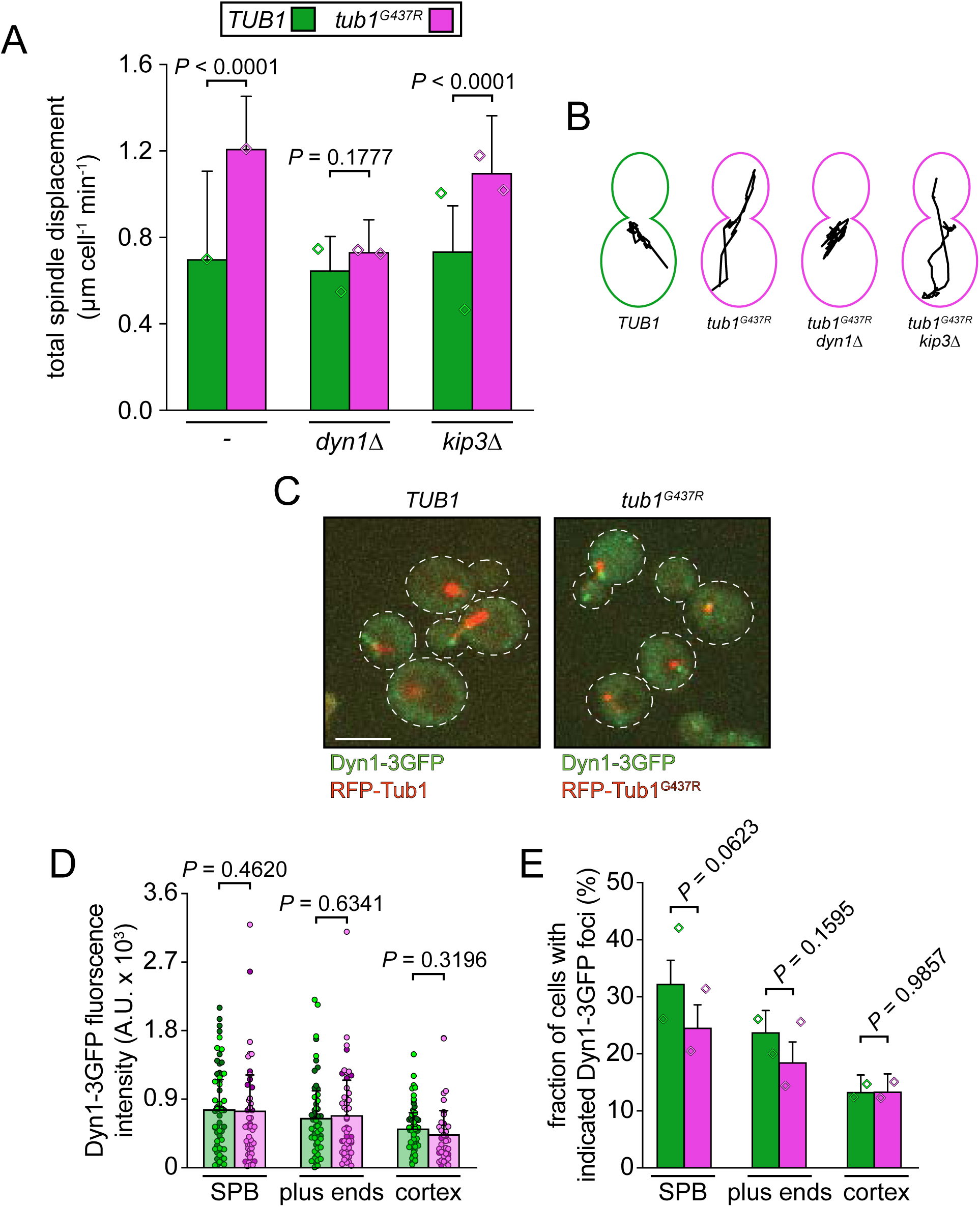
Enhanced spindle movement phenotype in *tub1*^*G437R*^ cells is dynein-dependent. (A) Plot depicting the average total displacement of the mitotic spindle in cells with the indicated genotype per minute of the acquisition period (for strains from left to right, n = 63, 24, 35, 35, 55, and 35 cells from two independent clones, respectively; *P* values were calculated using an unpaired Welch’s t test). (B) Representative displacement traces of the spindle noted over the 10 min acquisition duration for the indicated strains. (C) Representative fluorescence images of cells expressing Dyn1-3GFP and RFP-Tub1 (wild-type or mutant, as indicated). Scale bar, 5 µm. (D and E) Fluorescence intensity measurements of (D) and fraction of cells with (E) Dyn1-3GFP foci at the indicated subcellular localization in *TUB1* and *tub1*^*G437R*^ strains (for D, data sets from left to right, n = 56, 52, 58, 49, 60, and 36 cells, from two independent replicates; values from independent replicates are shown in two shades of green and magenta; for E, n = 248 and 232 cells, for wild-type and mutant cells respectively, from two independent replicates). For panel A, *P* values were calculated using an unpaired Welch’s t test. Diamonds represent mean values obtained from each independent replicate experiment.

### G437R microtubules exhibit reduced interaction with the dynein regulator She1

We wondered whether the apparent increase in dynein activity in *tub1*^*G437R*^ cells could be a consequence of reduced microtubule binding by the microtubule associated protein (MAP) She1, a dynein inhibitor (Markus et al., 2012; Markus et al., 2011; Woodruff et al., 2009). Since She1 inhibitory activity requires its microtubule binding activity (Ecklund et al., 2017), we first tested the effect of G437R on the interaction between She1 and tubulin. A She1-Gal4 activation domain (AD) fusion was tested for a two-hybrid interaction with either Tub1 or Tub1^G437R^, the latter of which were fused to the LexA DNA-binding domain (LexA_DBD_). Bim1, which is known to interact with α-tubulin in a two-hybrid assay (Krogan et al., 2006; Schwartz et al., 1997) was used as positive control, as was the kinesin Kip3. As expected, Bim1, Kip3 and She1 all showed a two-hybrid interaction with the Tub1 bait (Fig. 5A; positive interactions are apparent by growth on media lacking histidine, “-HIS”). Interestingly, the interaction between She1 and Tub1 – but not between Tub1 and either Bim1 or Kip3 – was reduced to background levels by the G437R mutation (Fig. 5A).

**Figure 5.**
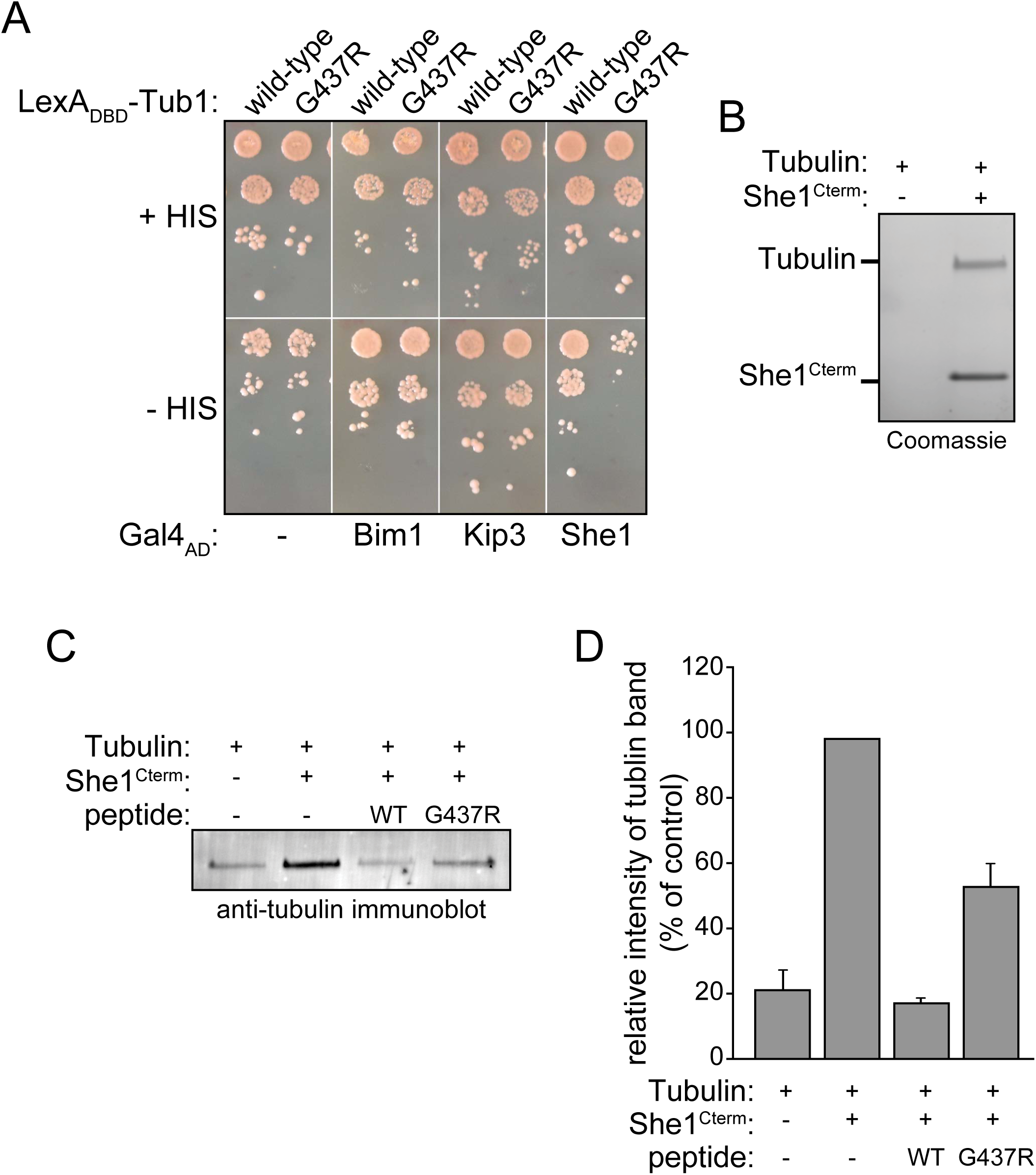
She1-Tubulin binding is reduced by the G437R mutation. (A) Two-hybrid assay illustrating the relative degree of interaction between tubulin (wild-type or mutant) and either the microtubule end binding protein Bim1 (homolog of human EB1), the kinesin-8 Kip3, or the dynein inhibitor She1. Serial dilutions of cells were spotted on minimal sold media with (top) or without histidine (bottom). Growth on histidine-deficient media is indicative of an interaction between bait and prey proteins. (B) Coomassie-stained acrylamide gel illustrating representative pull-down of cow brain tubulin by NiNTA-immobilized 6His-She1^Cter^. (C and D) A representative anti-tubulin immunoblot (C) and quantitation (D) of an experiment in which a peptide corresponding to the tubulin C-terminal tail (wild-type or mutant) was used to compete tubulin binding off of a NiNTA-immobilized 6His-She1^Cter^ (n = 2 independent replicates).

Like many MAPs, the interaction between She1 and microtubules requires the disordered C-terminal tails of tubulin (Ecklund et al., 2017; Markus et al., 2012). Thus, to further confirm the importance of G437 in the She1-tubulin interaction, we performed a pull-down assay in which this interaction was competitively inhibited by addition of a peptide encompassing the C-terminus of Tub1 (amino acids 415-447; both She1 and tubulin were used at 5 µg/ml or below, well below the critical concentration required for microtubule assembly). To this end, a 6His-She1 C-terminal fragment (She1^Cterm^; residues 194 to 338; which is sufficient for microtubule binding (Zhu et al., 2017)) was incubated with tubulin (see Fig. 5B for specificity of tubulin-She1^Cterm^ interaction) in the absence or presence of a peptide corresponding to the C-terminus of either wild-type or G437R tubulin (see Methods). The relative extent to which the NiNTA-bead-immobilized She1^Cterm^ pulled down tubulin was subsequently assessed by SDS-PAGE and immunoblot (using an antibody against a-tubulin). This revealed that the She1^Cterm^-tubulin interaction was strongly competed by the wild-type peptide, but much less so by the same peptide with the G437R mutation (Fig. 5C and D).

Next, we measured the extent to which She1 localizes to microtubules in cells by comparing the relative recruitment of full-length She1 to spindle microtubules (where She1 fluorescence is most prominent) in either wild-type or *tub1*^*G437R*^ cells. Consistent with the two-hybrid and *in vitro* data described above, we found that spindle-localized She1 fluorescence was 55.7% lower in G437R mutant cells than in wild-type cells (Fig. 6A and B). Interestingly, we also noted that the relative fluorescence intensity of mRuby2-Tub1^G437R^ in *tub1*^*G437R*^ cells was reduced to a similar extent (by 41.9%) with respect to mRuby2-Tub1 in wild-type cells (Fig. 6A and B). We reasoned this was due to one of two possible scenarios: (1) that the number of microtubules in the mitotic spindle is reduced as a consequence of the G437R mutation, or (2) that the wild-type copy of Tub3 – the less prevalent α-tubulin in budding yeast (see above) – is compensating for a reduced capacity of Tub1^G437R^ to incorporate into spindle microtubules. Since a reduced overall microtubule mass (scenario 1 above) would likely lead to severe consequences (*e.g.*, chromosome segregation defects, reduced viability) that were not apparent by live cell imaging (see Fig. 1C) or apparent cell viability (see Fig. 1E, “YPAD”), we focused on the latter possibility. To this end, we measured the spindle-localized fluorescence intensity of mRuby2-Tub3 in either wild-type or *tub1*^*G437R*^ cells and found a 92.3% increase in the extent to which Tub3 is incorporated into the spindles of mutant cells (Fig. 6A and B). Thus, the reduction in Tub1^G437R^ incorporation into spindle microtubules is entirely offset by a compensatory incorporation of Tub3 (Fig. 6D). These data indicate that the Tub1^G437R^ mutant tubulin is less able to incorporate into microtubules, and also explains the reliance of *tub1*^*G437R*^ mutant cells on *TUB3* (see Fig. 1D).

**Figure 6.**
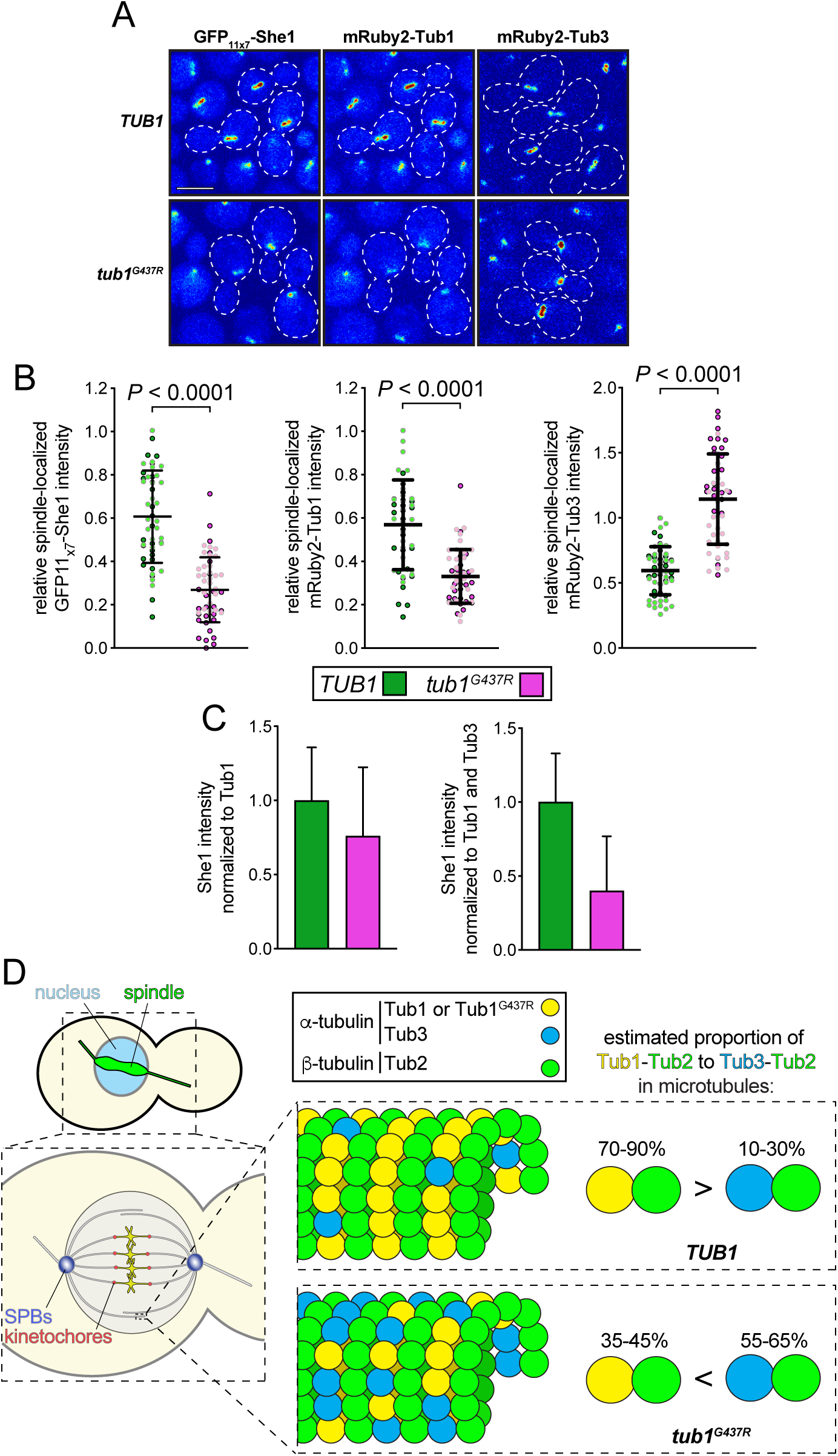
*In vivo* microtubule-binding by She1 is reduced by the G437R mutation. (A) Representative fluorescence images (shown as a heat map) depicting the relative degree of spindle-localized She1, Tub1 (wild-type or mutant), or Tub3 in wild-type or mutant cells. Scale bar, 4 µm. (B) Quantification of indicated spindle-localized molecule in wild-type or mutant cells. Prior to imaging, cells were arrested with 200 mM hydroxyurea (HU) for 2.5 hours (to enrich for cells with mitotic spindles), and then mounted on agarose pads containing HU for fluorescence microscopy (for datasets from left to right, n = 52, 52, 52, 52, 59, and 53 cells from two independent replicates, respectively; *P* values were calculated using an unpaired Welch’s t test). Values from independent replicates are shown in two shades of green and magenta. Bars indicate mean and standard deviation. (C) She1 intensity values normalized and corrected for relative Tub1 intensity (left), or Tub1 and Tub3 intensities (right; see Methods). (D) Cartoon depicting the relative shift in the balance of Tub1-Tub2 heterodimer:Tub3-Tub2 heterodimer from wild-type to mutant cells. Previous studies estimated that Tub3 accounts for roughly ∼10-30% of the cell’s α-tubulin content (Bode et al., 2003; Gartz Hanson et al., 2016). From the respective 42% reduction and 92% increase in mRuby2-Tub1 and mRuby3-Tub3 fluorescence intensities in the mutant cells, we estimate that this relative abundance is shifted such that Tub3 now accounts for roughly 55-65% of the cell’s α-tubulin content (although we only measured the relative abundance of each tubulin in spindle microtubules, this shift likely applies to both spindle and astral microtubules).

Finally, we used the relative Tub1/Tub3 spindle intensity values to calculate corrected spindle-localized intensity values for She1. The purpose of this correction was to normalize the degree of apparent She1-spindle binding to the relative microtubule mass. For instance, when She1 intensity values were normalized to Tub1 intensities – both of which were reduced in the mutant cells – the relative degree of She1-microtubule binding was only minimally reduced (Fig. 6C, left). However, when these normalized values were further corrected to account for the increased spindle incorporation of Tub3 into the spindles in the mutant cells, this revealed that She1 binding to Tub1^G437R^-containing microtubules is reduced by approximately 60% (Fig. 6C, right). Given the fact that the microtubules in the mutant cells are comprised of a roughly equal mixture of Tub1^G437R^ and Tub3 (see Fig. 6D, and Discussion), the reduced spindle intensity of She1 in the mutant cells is likely underestimating the degree by which the mutant tubulin reduces She1-microtubule binding. Taken together, these findings indicate that She1 binding to Tub1^G437R^-containing microtubules is severely compromised, thus accounting for the hyperactive dynein phenotype in these cells.

## Discussion

In summary, we have used budding yeast to characterize the consequences of the G437R α-tubulin mutation (equivalent to G436R in TUBA1A), which is likely causative of MCD in a human patient. Our results indicate that this mutation retains the ability to assemble into microtubules, albeit to a lesser extent than wild-type α-tubulin, which is apparent by the reduction of mRuby2-Tub1^G437R^ fluorescence in spindle microtubules. The reduced incorporation of the mutant tubulin into microtubules appears to be compensated by a corresponding increase in Tub3 incorporation in the *tub1*^*G437R*^ mutant, and explains the reliance of *tub1*^*G437R*^ mutant cells on Tub3 expression. Previous studies estimated that Tub3 accounts for roughly ∼10-30% of the cell’s α-tubulin content (Bode et al., 2003; Gartz Hanson et al., 2016). From the respective 42% reduction and 92% increase in Tub1^G437R^ and Tub3 microtubule incorporation in the mutant cells, we estimate that the relative abundance of Tub3:Tub1 in mutant microtubules is shifted such that Tub3 now accounts for 55-65% of the cell’s α-tubulin content (Fig. 6D). It is interesting to note that one of these studies (Gartz Hanson et al., 2016) observed a ∼17% increase in the rate of microtubule polymerization that was dependent on the presence of a wild-type copy of Tub3. This suggests that the increased proportion of Tub3 in the mutant microtubules may be partially responsible for the altered dynamicity observed here (16% increase in polymerization rate; Fig. 2).

It is interesting to note that the effects of G437R on dynamic instability were only significant during the G2/M phase of the cell cycle, at which point the growth and shrinkage rates, and the overall dynamicity increase significantly as a consequence of the mutation. Although the reasons for the cell cycle-dependent differences are unclear, they may be due to an inability (or increased ability) of a G2/M-specific factor to bind to and affect microtubule dynamics (either from the plus end, or along the lattice). Given the importance of the C-terminal tail in microtubule binding by numerous factors, and the proximity of G437 to the C-terminal tail of α-tubulin (see Fig. 1A and B), the possibility that this mutation is in fact impacting the structure/function of this region is a likely scenario (see Figs. S1 and S2). Although it is unclear how She1 affects microtubule dynamics, it’s reduced binding affinity for the mutant microtubules raises the possibility that the altered dynamics are a direct consequence of disrupted She1-microtubule binding, or an indirect cause of She1’s inability to modulate dynein or dynein-dynactin (see below).

The most notable phenotype in *tub1*^*G437R*^ cells was the dramatic increase in spindle translocation events throughout the mother and daughter cells (see Fig. 3 and Video S1). Spindle movements in budding yeast occur coincidentally with nuclear movement (due to the closed mitosis that takes place in this organism) and involves (1) alignment of the spindle along the mother-bud axis (a Kar9/actomyosin-mediated process), and (2) nuclear migration toward, and into the bud (a dynein-mediated process). Motor proteins are key effectors of this process since they can directly modulate microtubule dynamics (*e.g.*, Kip2, Kip3), and generate pushing and pulling forces via astral microtubules (Carvalho et al., 2004; Chen et al., 2019; Fukuda et al., 2014). In *tub1*^*G437R*^ cells, orientation of the mitotic spindle along the mother-bud axis (by the Kar9/actomyosin pathway) was not apparently disrupted (see Fig. S4), whereas dynein-mediated microtubule pulling was dramatically enhanced. Several lines of evidence point to She1 as the main molecular effector of this phenotype. To begin with, G437 is situated on a region of the microtubule surface that is proximal to known contact points for several MAPs (*e.g.*, Tau, Tpx2; see Fig. S1). Our data demonstrate that She1’s interaction with mutated microtubules or tubulin is impaired *in vivo*, and in two hybrid and pull-down assays. Finally, *tub1*^*G437R*^ cells exhibit an increase in dynein-mediated spindle movements in a manner similar to what has been observed in *she1Δ* cells (Markus et al., 2012). She1 exhibits the unique ability to specifically regulate dynein by reducing dynein’s microtubule dissociation rate, and consequently reducing its rate of motility (Ecklund et al., 2017). Therefore, it is logical that reducing the affinity of She1 for microtubules (as noted in *tub1*^*G437R*^ cells) would lead to an enhancement of dynein-mediated pulling forces similar to what was observed in *she1Δ* cells.

Also of interest is the apparent change in microtubule dynamics we observed specifically during and subsequent to dynein-mediated microtubule sliding events (see Fig. 3E). This decreased frequency of catastrophe events resulted in a greater proportion of long microtubules in the *tub1*^*G437R*^ strain, which may also partly account for the increase in dynein activity. It has been shown that increased microtubule lengths directly correlate with enhanced dynein activity in cells (Estrem et al., 2017). Although the role of She1 in affecting microtubule dynamics is not clear, it is possible that dynein-mediated depolymerizing activity – as has been noted *in vivo* (Estrem et al., 2017) – requires microtubule-bound She1. In particular, Estrem *et al*. (2017) recently showed that microtubules undergo a catastrophe event coincident with a dynein-mediated sliding event. They proposed that offloading of dynein-dynactin (the latter of which is a critical regulator of dynein activity) from microtubule plus ends to the cell cortex shifts the balance such that dynactin – which presumably stabilizes microtubules – is depleted from plus ends, while sufficient levels of dynein – which destabilizes microtubules – remain plus end-associated. In addition, previous studies found that She1 plays a role in precluding dynein-dynactin interaction at microtubule plus ends (Markus et al., 2011; Woodruff et al., 2009). Thus, either dynein-mediated microtubule destabilization or dynactin-mediated microtubule stabilization might be enhanced or reduced, respectively, by microtubule-bound She1. Although it is unclear if She1 needs to bind microtubules to affect dynein-dynactin interaction, it is possible that the Tub1^G437R^-mediated reduction in She1-microtubule binding might also enhance dynein-mediated recruitment of dynactin to plus ends, which would presumably provide a microtubule stabilization effect (due to the increased presence of dynactin). These models are not mutually exclusive, and may in fact both be acting to affect microtubule length during dynein-mediated spindle movement events.

Given the consequences on apparent brain development in the patient with TUBA1A^G436R^ (pachygyria, and severe microcephaly associated with postural delay and poor communication abilities), and the strong link between mutations in dynein activity and motor neuron diseases and developmental brain disorders (Bahi-Buisson et al., 2014; Laquerriere et al., 2017; Marzo et al., 2019; Vissers et al., 2010; Willemsen et al., 2012), our data linking disrupted dynein activity with this mutation are not entirely surprising. For example, dynein activity is critical for various aspects of early neuronal development, in part by promoting interkinetic nuclear migration in neuronal progenitors, and in the subsequent migration of the resulting postmitotic neurons (Del Bene et al., 2008; Hu et al., 2013; Tsai et al., 2010). Moreover, by effecting retrograde transport in neurons throughout their developmental progression, dynein activity is crucial for the maintenance of neuronal health, especially in motor neurons, in which cargoes must be transported over very long distances (≤ 1 m) (Bowman et al., 2000; Fu and Holzbaur, 2013; He et al., 2005; Hendricks et al., 2010; Rao et al., 2017; Shah et al., 2000; Wagner et al., 2004). However, of note, our findings indicate that dynein itself is unaffected by the mutation; rather, dynein activity is indirectly affected by the reduced microtubule binding affinity of a key regulatory MAP, She1. To date, a clear functional homolog of She1 in humans has not been identified. However, there indeed exists a myriad and complex network of MAPs in higher eukaryotes that may play similar roles to She1. For instance, the mammalian tau-related MAP4 protein, which binds in close proximity to G436 (Shigematsu et al., 2018), has been implicated in the control of dynein-mediated spindle orientation during mitosis in mammalian cells (Samora et al., 2011). MAP4 was also shown to physically interact with dynein-dynactin *in vivo* and to inhibit dynein-mediated microtubule gliding *in vitro* (Samora et al., 2011). MAP4 has also been shown to shorten dynein-dependent runs of melanosomes in *Xenopus* melanophores (Semenova et al., 2014). Another potential functional homolog of She1 is MAP9 (also known as ASter-Associated Protein, or ASAP), depletion of which disrupts spindle organization (Saffin et al., 2005; Venoux et al., 2008) and was recently shown to inhibit processive motility of purified dynein-dynactin complexes by specifically precluding microtubule binding by dynactin (Monroy et al., 2020). Thus, it will be important to determine how TUBA1A^G436R^ affects binding of these important neuronal MAPs.

## Material and methods

### Plasmids, yeast strain growth, and genetic manipulation

Strains used in this study were isogenic to either BY4742 (for Figure 1C and E, Figure 2, and Figure 3; *MAT*α; *ura3*Δ0 *leu2*Δ0 *his3*Δ1 *lys2*Δ0; provided by euroscarf http://www.euroscarf.de), or YEF473 (for Figure 1D, and Figure 6; *ura3-52 lys2-801 leu2-Δ1 his3-Δ200 trp1-Δ63*). The *TUB1* integrating plasmid, pCR2-TUB1 consists of the region of the *TUB1* locus from the intron (situated close the 5’ end of the gene) to 385 bp after the stop codon cloned into the pCR2 vector (Invitrogen). The *HIS3* gene expression cassette was ligated into the BsrGI site within the 3’ untranslated region of the *TUB1* sequence within pCR2 (pCR2-*TUB1*). The G437R mutation was subsequently introduced into pCR2-*TUB1* by PCR, generating pCR2-*tub1*^*G437R*^. For integration into the native *TUB1* locus, pCR2-*TUB1* (either wild-type or mutant) was digested with SphI, transformed into yeast using the lithium acetate method, and transformants were selected on media lacking histidine. All transformants were confirmed by PCR and sequencing.

RFP-TUB1 is derived from pAF125 as previously described (Caudron et al. 2008). In addition to this construct, we also generated pHIS3p:*mRuby2-tub1*^*G437R*^*+3’UTR*::*LEU2* to visualize microtubules in mutant cells. To this end, we engineered the G437R point mutation into pHIS3p:*mRuby2*-*TUB1+3’UTR*::*LEU2* (Markus et al., 2015) using traditional molecular biological methods. For comparison of relative a-tubulin incorporation into mitotic spindles, we used yeast strains with similarly integrated mRuby2-a-tubulins (pHIS3p:mRuby2-*TUB1+3’UTR*::*LEU2*, or pHIS3p:*mRuby2*-tub1^G437R^+3’UTR::*LEU2*). To assess relative incorporation of Tub3 into the mitotic spindle, we replaced the *TUB1+3’UTR* cassette in pHIS3p:*mRuby2*-*TUB1+3’UTR*::*TRP1* (Markus et al., 2015) with the *TUB3* genomic sequence, including 150 bp of the 3’UTR. This plasmid, pHIS3p:*mRuby2*-*TUB3+3’UTR*::*TRP1*, was digested with BbvCI, transformed into yeast using the lithium acetate method, and transformants were selected on media lacking tryptophan. pGFP-Bik1 (Lin et al., 2001) was kindly provided by D. Pellman.

For the two-hybrid experiments, the *TUB1* or *tub1*^*G437R*^ open reading frames from the respective pCR2 vectors (see above) were cloned into the pLexA vector (Addgene) to produce LexA_DBD_-Tub1 (or LexA_DBD_-Tub1^G437R^) fusion proteins with a-tubulin and the DNA-binding domain of LexA (LexA_DBD_). Bim1, Kip3, and She1 were amplified from genomic DNA and cloned into the pGADT7 vector (Invitrogen) to produce fusion proteins with the GAL4 activating domain (GAL4_AD_).

The 6His-tagged-She1^Cter^ expression plasmid for was constructed in pet28 vector (Novagen). The She1 C-terminal part coding for amino-acid 194 to 338 was cloned between the NdeI and XhoI site of the vector downstream of 6His and thrombin site.

To generate a GFP11_x7_-She1-expressing yeast strain (She1 fused to 7 copies of strand 11 of GFP) (Kamiyama et al., 2016), the GFP11_x7_ cassette was PCR amplified from pACUH:GFP11×7-mCherry-beta-tubulin (Addgene, plasmid # 70218), and integrated at the 5’ end of the *SHE1* open reading frame using the sequential *URA3* selection/5-FOA counterselection method. To separately express strands 1 thru 10 of the GFP barrel (GFP_1-10_, which is required to reconstitute fluorescence), we generated a plasmid with an expression cassette encoding GFP_1-10_ under the control of the strong *TEF1* promoter (*TEF1p*), as well as the *TRP1* selectable marker (pRS304:TEF1p:GFP_1-10_). We PCR amplified the *TEF1p:GFP1-10::TRP1* cassette and integrated it into the *lys2-801* locus using homologous recombination into the GFP_11×7_-She1-expressing yeast strain.

To assess viability on solid media (Figure 1E), serial dilutions of fresh overnight cultures of wild-type or *tub1*^*G437R*^ cells (three different haploid clones for each strain) were spotted onto solid YPAD media with or without benomyl, as indicated. Plates were incubated for two days at 30°C.

### Image acquisition and analysis

Cell imaging was performed on either a Zeiss Axiovert microscope equipped with a Cool Snap ES CCD camera (Ropper Scientific; for Figures 1 thru 4), or Nikon Ti-E microscope equipped with a 1.49 NA 100X TIRF objective, a Ti-S-E motorized stage, piezo Z-control (Physik Instrumente), an iXon DU888 cooled EM-CCD camera (Andor), a stage-top incubation system (Okolab), and a spinning disc confocal scanner unit (CSUX1; Yokogawa) with an emission filter wheel (ET525/50M for GFP, and ET632/60M for mRuby2; Chroma; Figure 6). For Figures 1 thru 4, 11 Z-planes with 0.3 μm spacing were captured using 2×2 binning (the exposure time varied between experiments). For Figure 6, 11 Z-planes with 0.3 μm spacing were captured.

For microtubule dynamics measurements, maximum intensity projection images of Bik1-GFP-expressing cells were used. Microtubule lengths at each time point were measured manually from maximum intensity projections, and microtubule dynamic parameters were calculated as described (Kosco et al., 2001), using an in-house Visual Basic macro in Excel (Caudron et al., 2008). We measured background-corrected She1, Tub1, or Tub3 spindle-localized fluorescence from maximum intensity projections using ImageJ (NIH). Tub1-corrected She1 intensities were calculated by normalizing the fluorescence intensity values of each (She1, Tub1 and Tub3) to 1 (by dividing the raw mean background corrected values for that measured in wild-type or *tub1*^*G437R*^ cells by the wild-type value), and then dividing the resulting normalized She1 values by the normalized Tub1 value for wild-type and *tub1*^*G437R*^, respectively. To additionally correct for Tub3 intensities, these Tub1-corrected values were then divided by the normalized Tub3 values (as calculated above; in which wild-type = 1, and *tub1*^*G437R*^ = 1.91).

To assess spindle dynamics parameters, we determined spindle position in cells over time by clicking the center of a preanaphase spindle in each frame, and calculating the displacement between frames using an in-house developed ImageJ macro. The spindle position with respect to the cell boundaries (Figure 3G) was determined for spindles with a pole-to-pole length of 0.8–1.2 μm from the first frames of movies of G2/M cells. The quantification was done blind to the genotype. At least 52 preanaphase spindles from 3 independent clones were measured for each strain, as described (DeZwaan et al., 1997).

### Production of recombinant protein

The 6His-She1^Cter^ plasmid was transformed into Rosetta pLysS bacteria. For protein production, bacterial cultures grown in M9 medium were induced by addition of 1 mM IPTG overnight at 18°C. The clarified lysate was separated on a cation exchange column (SP GE healthcare), and bound protein was eluted in cation elution buffer (500 mM NaCl, 40 mM Tris pH 9.2), and then applied to a NiNTA column (Qiagen) after addition of 20 mM imidazole. After elution in NiNTA elution buffer (300 mM NaCl, 20mM Tris pH 7.5, 250mM imidazole), the protein was simultaneously concentrated and buffer-exchanged (with a Vivaspin Turbo 4; Sartorius) into PM buffer (80 mM Pipes pH 6.8, 1mM MgCl2, 0.2% NP40).

### Pull-down experiments

For She1^Cterm^-tubulin pull-down experiments, 2.5 µg of purified tubulin (from cow brain) was incubated with or without 2.5 µg of 6His-She1^Cter^ in 500 µl of PM buffer. After 30 minutes at room temperature, 10 µl of NiNTA beads were added (Thermo Fisher). After a 10 minute incubation, the beads were washed three times with PM buffer, and then boiled for 5 minutes in 60 µl of sample buffer before SDS-PAGE analysis. For peptide competition experiments, 50 µg of each peptide (dissolved in PM buffer; wild-type : EEGEF TEARE DLAAL ERDYI EV**<u>G</u>**AD SYAEE EEF; G437R: EEGEF TEARE DLAAL ERDYI EV**<u>R</u>**AD SYAEE EEF) were added to the mixture of She1 (0.25 µg) and tubulin (1.25 µg) and allowed to bind for 30 min at room temperature. Then, 10 µl of NiNTA beads were added (Thermo Fisher), incubated for 10 minutes, washed three times with PM buffer, and then boiled for 5 minutes in 60 µl of sample buffer prior to SDS-PAGE and immunoblot blot analysis. For immunblotting, the gel was transferred to a PVDF membrane (BioRad TransBlot Turbo), blocked with PBS supplemented with 5% milk for 30 minutes, and then probed with anti-α-tubulin (YL1/2 1:10,000) followed by Cy5 anti-rat secondary antibody (Jackson ImmunoResearch, 1:2000). To quantify the tubulin signals, the local background was subtracted and signals were normalized such that tubulin-bound 6His-She1^Cter^ without peptide was equal to 100. The average of two replicates is presented in the graph (Figure 5D).

### Structural analyses and mutation modeling

Molecular graphics and analyses were performed with the UCSF Chimera package. The rotamers function was used to model an arginine substitution into position 436 of the α-tubulin structure (pdb 3J6G). The top 4 rotamers were selected for visualization (see Fig. S2). Chimera is developed by the Resource for Biocomputing, Visualization, and Informatics at the University of California, San Francisco (supported by NIGMS P41-GM103311).

## Acknowledgements

We would like to thank Jeff Moore for kindly providing a *tub3Δ* yeast strain. This work was funded by IINCA (TetraTips PLBIO10-030 to A.A. and C.B.) and the NIH/NIGMS (GM118492 to S.M.M.).

## Literature Cited

Adames, N.R., and J.A. Cooper. 2000. Microtubule interactions with the cell cortex causing nuclear movements in Saccharomyces cerevisiae. J Cell Biol. 149:863–874.

Alushin, G.M., G.C. Lander, E.H. Kellogg, R. Zhang, D. Baker, and E. Nogales. 2014. High-resolution microtubule structures reveal the structural transitions in alphabeta-tubulin upon GTP hydrolysis. Cell. 157:1117–1129.

Badin-Larcon, A.C., C. Boscheron, J.M. Soleilhac, M. Piel, C. Mann, E. Denarier, A. Fourest-Lieuvin, L. Lafanechere, M. Bornens, and D. Job. 2004. Suppression of nuclear oscillations in Saccharomyces cerevisiae expressing Glu tubulin. Proc Natl Acad Sci U S A. 101:5577–5582.

Bahi-Buisson, N., K. Poirier, N. Boddaert, Y. Saillour, L. Castelnau, N. Philip, G. Buyse, L. Villard, S. Joriot, S. Marret, M. Bourgeois, H. Van Esch, L. Lagae, J. Amiel, L. Hertz-Pannier, A. Roubertie, F. Rivier, J.M. Pinard, C. Beldjord, and J. Chelly. 2008. Refinement of cortical dysgeneses spectrum associated with TUBA1A mutations. Journal of medical genetics. 45:647–653.

Bahi-Buisson, N., K. Poirier, F. Fourniol, Y. Saillour, S. Valence, N. Lebrun, M. Hully, C.F. Bianco, N. Boddaert, C. Elie, K. Lascelles, I. Souville, L.I.-T. Consortium, C. Beldjord, and J. Chelly. 2014. The wide spectrum of tubulinopathies: what are the key features for the diagnosis? Brain. 137:1676–1700.

Bershteyn, M., T.J. Nowakowski, A.A. Pollen, E. Di Lullo, A. Nene, A. Wynshaw-Boris, and A.R. Kriegstein. 2017. Human iPSC-Derived Cerebral Organoids Model Cellular Features of Lissencephaly and Reveal Prolonged Mitosis of Outer Radial Glia. Cell Stem Cell. 20:435–449 e434.

Bode, C.J., M.L. Gupta, K.A. Suprenant, and R.H. Himes. 2003. The two alpha-tubulin isotypes in budding yeast have opposing effects on microtubule dynamics in vitro. EMBO Rep. 4:94–99.

Bowman, A.B., A. Kamal, B.W. Ritchings, A.V. Philp, M. McGrail, J.G. Gindhart, and L.S. Goldstein. 2000. Kinesin-dependent axonal transport is mediated by the sunday driver (SYD) protein. Cell. 103:583–594.

Cabernard, C., and C.Q. Doe. 2009. Apical/basal spindle orientation is required for neuroblast homeostasis and neuronal differentiation in Drosophila. Dev Cell. 17:134–141.

Carminati, J.L., and T. Stearns. 1997. Microtubules orient the mitotic spindle in yeast through dynein-dependent interactions with the cell cortex. J Cell Biol. 138:629–641.

Carvalho, P., M.L. Gupta, Jr., M.A. Hoyt, and D. Pellman. 2004. Cell cycle control of kinesin-mediated transport of Bik1 (CLIP-170) regulates microtubule stability and dynein activation. Dev Cell. 6:815–829.

Caudron, F., A. Andrieux, D. Job, and C. Boscheron. 2008. A new role for kinesin-directed transport of Bik1p (CLIP-170) in Saccharomyces cerevisiae. J Cell Sci. 121:1506–1513.

Chen, X., L.A. Widmer, M.M. Stangier, M.O. Steinmetz, J. Stelling, and Y. Barral. 2019. Remote control of microtubule plus-end dynamics and function from the minus-end. Elife. 8.

Das, R.M., and K.G. Storey. 2012. Mitotic spindle orientation can direct cell fate and bias Notch activity in chick neural tube. EMBO Rep. 13:448–454.

Del Bene, F., A.M. Wehman, B.A. Link, and H. Baier. 2008. Regulation of neurogenesis by interkinetic nuclear migration through an apical-basal notch gradient. Cell. 134:1055–1065.

Denarier, E., C. Brousse, A. Sissoko, A. Andrieux, and C. Boscheron. 2019. A neurodevelopmental TUBB2B beta-tubulin mutation impairs Bim1 (yeast EB1)- dependent spindle positioning. Biol Open. 8.

DeZwaan, T.M., E. Ellingson, D. Pellman, and D.M. Roof. 1997. Kinesin-related KIP3 of Saccharomyces cerevisiae is required for a distinct step in nuclear migration. J Cell Biol. 138:1023–1040.

Ecklund, K.H., T. Morisaki, L.G. Lammers, M.G. Marzo, T.J. Stasevich, and S.M. Markus. 2017. She1 affects dynein through direct interactions with the microtubule and the dynein microtubule-binding domain. Nat Commun. 8:2151.

Estrem, C., C.P. Fees, and J.K. Moore. 2017. Dynein is regulated by the stability of its microtubule track. J Cell Biol. 216:2047–2058.

Findeisen, P., S. Muhlhausen, S. Dempewolf, J. Hertzog, A. Zietlow, T. Carlomagno, and M. Kollmar. 2014. Six subgroups and extensive recent duplications characterize the evolution of the eukaryotic tubulin protein family. Genome Biol Evol. 6:2274–2288.

Fu, M.M., and E.L. Holzbaur. 2013. JIP1 regulates the directionality of APP axonal transport by coordinating kinesin and dynein motors. J Cell Biol. 202:495–508.

Fukuda, Y., A. Luchniak, E.R. Murphy, and M.L. Gupta, Jr. 2014. Spatial control of microtubule length and lifetime by opposing stabilizing and destabilizing functions of Kinesin-8. Curr Biol. 24:1826–1835.

Gartz Hanson, M., J. Aiken, D.V. Sietsema, D. Sept, E.A. Bates, L. Niswander, and J.K. Moore. 2016. Novel alpha-tubulin mutation disrupts neural development and tubulin proteostasis. Dev Biol. 409:406–419.

Gigant, B., W. Wang, B. Dreier, Q. Jiang, L. Pecqueur, A. Pluckthun, C. Wang, and M. Knossow. 2013. Structure of a kinesin-tubulin complex and implications for kinesin motility. Nat Struct Mol Biol. 20:1001–1007.

Gupta, M.L., Jr., P. Carvalho, D.M. Roof, and D. Pellman. 2006. Plus end-specific depolymerase activity of Kip3, a kinesin-8 protein, explains its role in positioning the yeast mitotic spindle. Nat Cell Biol. 8:913–923.

He, Y., F. Francis, K.A. Myers, W. Yu, M.M. Black, and P.W. Baas. 2005. Role of cytoplasmic dynein in the axonal transport of microtubules and neurofilaments. J Cell Biol. 168:697–703.

Heil-Chapdelaine, R.A., J.R. Oberle, and J.A. Cooper. 2000. The cortical protein Num1p is essential for dynein-dependent interactions of microtubules with the cortex. J Cell Biol. 151:1337–1344.

Hendricks, A.G., E. Perlson, J.L. Ross, H.W. Schroeder, 3rd, M. Tokito, and E.L. Holzbaur. 2010. Motor coordination via a tug-of-war mechanism drives bidirectional vesicle transport. Curr Biol. 20:697–702.

Hu, D.J., A.D. Baffet, T. Nayak, A. Akhmanova, V. Doye, and R.B. Vallee. 2013. Dynein recruitment to nuclear pores activates apical nuclear migration and mitotic entry in brain progenitor cells. Cell. 154:1300–1313.

Hwang, E., J. Kusch, Y. Barral, and T.C. Huffaker. 2003. Spindle orientation in Saccharomyces cerevisiae depends on the transport of microtubule ends along polarized actin cables. J Cell Biol. 161:483–488.

Kamiyama, D., S. Sekine, B. Barsi-Rhyne, J. Hu, B. Chen, L.A. Gilbert, H. Ishikawa, M.D. Leonetti, W.F. Marshall, J.S. Weissman, and B. Huang. 2016. Versatile protein tagging in cells with split fluorescent protein. Nat Commun. 7:11046.

Kellogg, E.H., N.M.A. Hejab, S. Poepsel, K.H. Downing, F. DiMaio, and E. Nogales. 2018. Near-atomic model of microtubule-tau interactions. Science. 360:1242–1246.

Khodiyar, V.K., L.J. Maltais, B.J. Ruef, K.M. Sneddon, J.R. Smith, M. Shimoyama, F. Cabral, C. Dumontet, S.K. Dutcher, R.J. Harvey, L. Lafanechere, J.M. Murray, E. Nogales, D. Piquemal, F. Stanchi, S. Povey, and R.C. Lovering. 2007. A revised nomenclature for the human and rodent alpha-tubulin gene family. Genomics. 90:285–289.

Kosco, K.A., C.G. Pearson, P.S. Maddox, P.J. Wang, I.R. Adams, E.D. Salmon, K. Bloom, and T.C. Huffaker. 2001. Control of microtubule dynamics by Stu2p is essential for spindle orientation and metaphase chromosome alignment in yeast. Mol Biol Cell. 12:2870–2880.

Krogan, N.J., G. Cagney, H. Yu, G. Zhong, X. Guo, A. Ignatchenko, J. Li, S. Pu, N. Datta, A.P. Tikuisis, T. Punna, J.M. Peregrin-Alvarez, M. Shales, X. Zhang, M. Davey, M.D. Robinson, A. Paccanaro, J.E. Bray, A. Sheung, B. Beattie, D.P. Richards, V. Canadien, A. Lalev, F. Mena, P. Wong, A. Starostine, M.M. Canete, J. Vlasblom, S. Wu, C. Orsi, S.R. Collins, S. Chandran, R. Haw, J.J. Rilstone, K. Gandi, N.J. Thompson, G. Musso, P. St Onge, S. Ghanny, M.H. Lam, G. Butland, A.M. Altaf-Ul, S. Kanaya, A. Shilatifard, E. O’Shea, J.S. Weissman, C.J. Ingles, T.R. Hughes, J. Parkinson, M. Gerstein, S.J. Wodak, A. Emili, and J.F. Greenblatt. 2006. Global landscape of protein complexes in the yeast Saccharomyces cerevisiae. Nature. 440:637–643.

Kumar, R.A., D.T. Pilz, T.D. Babatz, T.D. Cushion, K. Harvey, M. Topf, L. Yates, S. Robb, G. Uyanik, G.M. Mancini, M.I. Rees, R.J. Harvey, and W.B. Dobyns. 2010. TUBA1A mutations cause wide spectrum lissencephaly (smooth brain) and suggest that multiple neuronal migration pathways converge on alpha tubulins. Hum Mol Genet. 19:2817–2827.

Lacey, S.E., S. He, S.H. Scheres, and A.P. Carter. 2019. Cryo-EM of dynein microtubule-binding domains shows how an axonemal dynein distorts the microtubule. Elife. 8.

Lammers, L.G., and S.M. Markus. 2015. The dynein cortical anchor Num1 activates dynein motility by relieving Pac1/LIS1-mediated inhibition. J Cell Biol. 211:309–322.

Laquerriere, A., C. Maillard, M. Cavallin, F. Chapon, F. Marguet, A. Molin, S. Sigaudy, M. Blouet, G. Benoist, C. Fernandez, K. Poirier, J. Chelly, S. Thomas, and N. Bahi-Buisson. 2017. Neuropathological Hallmarks of Brain Malformations in Extreme Phenotypes Related to DYNC1H1 Mutations. J Neuropathol Exp Neurol. 76:195–205.

Lee, L., J.S. Tirnauer, J. Li, S.C. Schuyler, J.Y. Liu, and D. Pellman. 2000. Positioning of the mitotic spindle by a cortical-microtubule capture mechanism. Science. 287:2260–2262.

Lee, W.L., J.R. Oberle, and J.A. Cooper. 2003. The role of the lissencephaly protein Pac1 during nuclear migration in budding yeast. J Cell Biol. 160:355–364.

Li, Y.Y., E. Yeh, T. Hays, and K. Bloom. 1993. Disruption of mitotic spindle orientation in a yeast dynein mutant. Proc Natl Acad Sci U S A. 90:10096–10100.

Liakopoulos, D., J. Kusch, S. Grava, J. Vogel, and Y. Barral. 2003. Asymmetric loading of Kar9 onto spindle poles and microtubules ensures proper spindle alignment. Cell. 112:561–574.

Lin, H., P. de Carvalho, D. Kho, C.Y. Tai, P. Pierre, G.R. Fink, and D. Pellman. 2001. Polyploids require Bik1 for kinetochore-microtubule attachment. J Cell Biol. 155:1173–1184.

Lowe, J., H. Li, K.H. Downing, and E. Nogales. 2001. Refined structure of alpha betatubulin at 3.5 A resolution. J Mol Biol. 313:1045–1057.

Markus, S.M., K.A. Kalutkiewicz, and W.L. Lee. 2012. She1-mediated inhibition of dynein motility along astral microtubules promotes polarized spindle movements. Curr Biol. 22:2221–2230.

Markus, S.M., and W.L. Lee. 2011. Regulated offloading of cytoplasmic dynein from microtubule plus ends to the cortex. Dev Cell. 20:639–651.

Markus, S.M., S. Omer, K. Baranowski, and W.L. Lee. 2015. Improved Plasmids for Fluorescent Protein Tagging of Microtubules in Saccharomyces cerevisiae. Traffic. 16:773–786.

Markus, S.M., K.M. Plevock, B.J. St Germain, J.J. Punch, C.W. Meaden, and W.L. Lee. 2011. Quantitative analysis of Pac1/LIS1-mediated dynein targeting: Implications for regulation of dynein activity in budding yeast. Cytoskeleton (Hoboken). 68:157–174.

Marzo, M.G., J.M. Griswold, and S.M. Markus. 2020. Pac1/LIS1 stabilizes an uninhibited conformation of dynein to coordinate its localization and activity. Nat Cell Biol:In press.

Marzo, M.G., J.M. Griswold, K.M. Ruff, R.E. Buchmeier, C.P. Fees, and S.M. Markus. 2019. Molecular basis for dyneinopathies reveals insight into dynein regulation and dysfunction. Elife. 8.

Miller, R.K., and M.D. Rose. 1998. Kar9p is a novel cortical protein required for cytoplasmic microtubule orientation in yeast. J Cell Biol. 140:377–390.

Mohamed M. Elshenawy, E.K., Sara Volz, Janina Baumbach, Simon L. Bullock, Ahmet Yildiz. 2020. Lis1 activates dynein motility by pairing it with dynactin. Nat Cell Biol: In press.

Monroy, B.Y., T.C. Tan, J.M. Oclaman, J.S. Han, S. Simo, S. Niwa, D.W. Nowakowski, R.J. McKenney, and K.M. Ori-McKenney. 2020. A Combinatorial MAP Code Dictates Polarized Microtubule Transport. Dev Cell.

Nogales, E., S.G. Wolf, and K.H. Downing. 1998. Structure of the alpha beta tubulin dimer by electron crystallography. Nature. 391:199–203.

Pettersen, E.F., T.D. Goddard, C.C. Huang, G.S. Couch, D.M. Greenblatt, E.C. Meng, and T.E. Ferrin. 2004. UCSF Chimera--a visualization system for exploratory research and analysis. J Comput Chem. 25:1605–1612.

Poirier, K., N. Lebrun, L. Broix, G. Tian, Y. Saillour, C. Boscheron, E. Parrini, S. Valence, B.S. Pierre, M. Oger, D. Lacombe, D. Genevieve, E. Fontana, F. Darra, C. Cances, M. Barth, D. Bonneau, B.D. Bernadina, S. N’Guyen, C. Gitiaux, P. Parent, V. des Portes, J.M. Pedespan, V. Legrez, L. Castelnau-Ptakine, P. Nitschke, T. Hieu, C. Masson, D. Zelenika, A. Andrieux, F. Francis, R. Guerrini, N.J. Cowan, N. Bahi-Buisson, and J. Chelly. 2013. Mutations in TUBG1, DYNC1H1, KIF5C and KIF2A cause malformations of cortical development and microcephaly. Nature genetics. 45:639–647.

Rao, A.N., A. Patil, M.M. Black, E.M. Craig, K.A. Myers, H.T. Yeung, and P.W. Baas. 2017. Cytoplasmic Dynein Transports Axonal Microtubules in a Polarity-Sorting Manner. Cell Rep. 19:2210–2219.

Richards, K.L., K.R. Anders, E. Nogales, K. Schwartz, K.H. Downing, and D. Botstein. 2000. Structure-function relationships in yeast tubulins. Mol Biol Cell. 11:1887–1903.

Saffin, J.M., M. Venoux, C. Prigent, J. Espeut, F. Poulat, D. Giorgi, A. Abrieu, and S. Rouquier. 2005. ASAP, a human microtubule-associated protein required for bipolar spindle assembly and cytokinesis. Proc Natl Acad Sci U S A. 102:11302–11307.

Samora, C.P., B. Mogessie, L. Conway, J.L. Ross, A. Straube, and A.D. McAinsh. 2011. MAP4 and CLASP1 operate as a safety mechanism to maintain a stable spindle position in mitosis. Nat Cell Biol. 13:1040–1050.

Schatz, P.J., L. Pillus, P. Grisafi, F. Solomon, and D. Botstein. 1986a. Two functional alpha-tubulin genes of the yeast Saccharomyces cerevisiae encode divergent proteins. Mol Cell Biol. 6:3711–3721.

Schatz, P.J., F. Solomon, and D. Botstein. 1986b. Genetically essential and nonessential alpha-tubulin genes specify functionally interchangeable proteins. Mol Cell Biol. 6:3722–3733.

Schmidt, H., R. Zalyte, L. Urnavicius, and A.P. Carter. 2015. Structure of human cytoplasmic dynein-2 primed for its power stroke. Nature. 518:435–438.

Schwartz, K., K. Richards, and D. Botstein. 1997. BIM1 encodes a microtubule-binding protein in yeast. Mol Biol Cell. 8:2677–2691.

Semenova, I., K. Ikeda, K. Resaul, P. Kraikivski, M. Aguiar, S. Gygi, I. Zaliapin, A. Cowan, and V. Rodionov. 2014. Regulation of microtubule-based transport by MAP4. Molecular biology of the cell. 25:3119–3132.

Shah, J.V., L.A. Flanagan, P.A. Janmey, and J.F. Leterrier. 2000. Bidirectional translocation of neurofilaments along microtubules mediated in part by dynein/dynactin. Mol Biol Cell. 11:3495–3508.

Sheeman, B., P. Carvalho, I. Sagot, J. Geiser, D. Kho, M.A. Hoyt, and D. Pellman. 2003. Determinants of S. cerevisiae dynein localization and activation: implications for the mechanism of spindle positioning. Curr Biol. 13:364–372.

Shigematsu, H., T. Imasaki, C. Doki, T. Sumi, M. Aoki, T. Uchikubo-Kamo, A. Sakamoto, K. Tokuraku, M. Shirouzu, and R. Nitta. 2018. Structural insight into microtubule stabilization and kinesin inhibition by Tau family MAPs. J Cell Biol. 217:4155–4163.

Tan, R., A.J. Lam, T. Tan, J. Han, D.W. Nowakowski, M. Vershinin, S. Simo, K.M. Ori-McKenney, and R.J. McKenney. 2019. Microtubules gate tau condensation to spatially regulate microtubule functions. Nat Cell Biol. 21:1078–1085.

Toso, R.J., M.A. Jordan, K.W. Farrell, B. Matsumoto, and L. Wilson. 1993. Kinetic stabilization of microtubule dynamic instability in vitro by vinblastine. Biochemistry. 32:1285–1293.

Tsai, J.W., Y. Chen, A.R. Kriegstein, and R.B. Vallee. 2005. LIS1 RNA interference blocks neural stem cell division, morphogenesis, and motility at multiple stages. J Cell Biol. 170:935–945.

Tsai, J.W., S. Lian Wn Fau - Kemal, A.R. Kemal S Fau - Kriegstein, R.B. Kriegstein Ar Fau - Vallee, and R.B. Vallee. 2010. Kinesin 3 and cytoplasmic dynein mediate interkinetic nuclear migration in neural stem cells. Nat Neuroscience. 13:1463–1471.

Venoux, M., J. Basbous, C. Berthenet, C. Prigent, A. Fernandez, N.J. Lamb, and S. Rouquier. 2008. ASAP is a novel substrate of the oncogenic mitotic kinase Aurora-A: phosphorylation on Ser625 is essential to spindle formation and mitosis. Hum Mol Genet. 17:215–224.

Vissers, L.E., J. de Ligt, C. Gilissen, I. Janssen, M. Steehouwer, P. de Vries, B. van Lier, P. Arts, N. Wieskamp, M. del Rosario, B.W. van Bon, A. Hoischen, B.B. de Vries, H.G. Brunner, and J.A. Veltman. 2010. A de novo paradigm for mental retardation. Nature genetics. 42:1109–1112.

Wagner, O.I., J. Ascano, M. Tokito, J.F. Leterrier, P.A. Janmey, and E.L. Holzbaur. 2004. The interaction of neurofilaments with the microtubule motor cytoplasmic dynein. Mol Biol Cell. 15:5092–5100.

Willemsen, M.H., L.E. Vissers, M.A. Willemsen, B.W. van Bon, T. Kroes, J. de Ligt, B.B. de Vries, J. Schoots, D. Lugtenberg, B.C. Hamel, H. van Bokhoven, H.G. Brunner, J.A. Veltman, and T. Kleefstra. 2012. Mutations in DYNC1H1 cause severe intellectual disability with neuronal migration defects. Journal of medical genetics. 49:179–183.

Williams, S.E., S. Beronja, H.A. Pasolli, and E. Fuchs. 2011. Asymmetric cell divisions promote Notch-dependent epidermal differentiation. Nature. 470:353–358.

Woodruff, J.B., D.G. Drubin, and G. Barnes. 2009. Dynein-driven mitotic spindle positioning restricted to anaphase by She1p inhibition of dynactin recruitment. Mol Biol Cell. 20:3003–3011.

Wu, M., C.L. Smith, J.A. Hall, I. Lee, K. Luby-Phelps, and M.D. Tallquist. 2010. Epicardial spindle orientation controls cell entry into the myocardium. Dev Cell. 19:114–125.

Yin, H., D. Pruyne, T.C. Huffaker, and A. Bretscher. 2000. Myosin V orientates the mitotic spindle in yeast. Nature. 406:1013–1015.

Yingling, J., Y.H. Youn, D. Darling, K. Toyo-Oka, T. Pramparo, S. Hirotsune, and A. Wynshaw-Boris. 2008. Neuroepithelial stem cell proliferation requires LIS1 for precise spindle orientation and symmetric division. Cell. 132:474–486.

Zaw Min Htet, J.P.G., Richard W. Baker, Andres E. Leschziner, Morgan E. DeSantis, Samara L. Reck-Peterson. 2020. Lis1 promotes the formation of activated cytoplasmic dynein-1 complexes. Nat Cell Biol:In press.

Zhang, R., J. Roostalu, T. Surrey, and E. Nogales. 2017. Structural insight into TPX2-stimulated microtubule assembly. Elife. 6.

Zhu, Y., X. An, A. Tomaszewski, P.K. Hepler, and W.L. Lee. 2017. Microtubule crosslinking activity of She1 ensures spindle stability for spindle positioning. J Cell Biol. 216:2759–2775.

